# Chemogenetic manipulation of endogenous proteins in fission yeast using a self-localizing ligand-induced protein translocation (SLIPT) system

**DOI:** 10.1101/2023.07.27.550911

**Authors:** Akinobu Nakamura, Yuhei Goto, Hironori Sugiyama, Shinya Tsukiji, Kazuhiro Aoki

## Abstract

Cells sense extracellular stimuli through membrane receptors and process the information through an intracellular signaling network. Protein translocation triggers intracellular signaling, and techniques such as chemically induced dimerization (CID) have been used to manipulate signaling pathways by altering the subcellular localization of signaling molecules. However, in the fission yeast *Schizosaccharomyces pombe*, the commonly used FKBP-FRB system has technical limitations, and therefore perturbation tools with low cytotoxicity and high temporal resolution are needed. We here applied our recently developed self-localizing ligand-induced protein translocation (SLIPT) system to *S. pombe* and successfully perturbed several cell cycle-related proteins. The SLIPT system utilizes self-localizing ligands to recruit binding partners to specific subcellular compartments, such as the plasma membrane or nucleus. We optimzed the self-localizing ligands to maintain long-term recruitment of target molecules to the plasma membrane. By knocking in genes encoding the binding partners for self-localizing ligands, we observed changes in the localization of several endogenous molecules and found perturbations in the cell cycle and associated phenotypes. This study demonstrates the effectiveness of the SLIPT system as a chemogenetic tool for rapid perturbation of endogenous molecules in *S. pombe*, providing a valuable approach for studying intracellular signaling and cell cycle regulation with improved temporal resolution.

## Introduction

Cells sense extracellular stimuli through their receptors on the plasma membrane (PM) and process the information through an intracellular signaling network ^1^. The intracellular signaling network consists of multiple biophysical diffusions and reactions, including protein-protein interactions and enzymatic reactions. It is essential to understand the mechanisms of intracellular signaling because several diseases, such as malignant tumors, arise from genetic dysfunction of intracellular signaling molecules ^2,3^. Although gene knockout has been widely used to study the physiological function of signaling proteins, it is difficult to analyze the gene when the gene knockout causes lethality. Furthermore, because the effects of gene knockout are long-lasting, the changes caused by gene knockout may be masked by adaptations in the signal transduction system ^4,5^. Therefore, tools are needed to rapidly control the activity of signaling proteins with high temporal resolution.

Protein translocation is known to trigger intracellular signaling. For example, Akt, a Ser/Thr protein kinase, is recruited to and activated at the PM by binding to phosphatidyl inositol trisphosphate PI(3,4,5)P3, leading to the initiation of downstream signaling ^6^. In addition, protein translocation restricts signaling to specific subcellular compartments. Upon growth factor stimulation, ERK, a classical MAP kinase, is phosphorylated and activated in the cytoplasm, followed by nuclear translocation to phosphorylate nuclear substrates such as transcription factors ^7^. Several techniques have been reported to rapidly control signaling pathways by altering the subcellular localization of intracellular signaling molecules. Among them, chemically induced dimerization (CID) has been widely used to manipulate various signaling pathways, such as the Ras-ERK MAP kinase pathway, the PI3K-Akt pathway, and the Rho family G protein pathway ^8–12^. CID is a chemical process in which two proteins bind only in the presence of small chemical dimerizers ^13^, and in many cases the signaling system is manipulated by inducing heterodimerization of the proteins. The most widely used CID system is the heterodimerization of FKBP-FRB by rapamycin ^8,14^. However, chemically induced heterodimerization requires the expression of two types of proteins, namely, a localizer and an actuator. Recently, we have developed a self-localizing ligand-induced protein translocation (SLIPT) system, which requires only an actuator and a small compound with a subcellular localization tag (self-localizing ligand). By using the SLIPT system, we have successfully altered the subcellular localization of actuators and manipulated the Grb2/SOS/Ras/ERK, PI3K, Tiam/Rac, Gqα/Ca^2+^, Gsα/cAMP, and PKCδ signaling pathways in mammalian cells ^15–22^ (Figure 1A).

**Figure 1.**
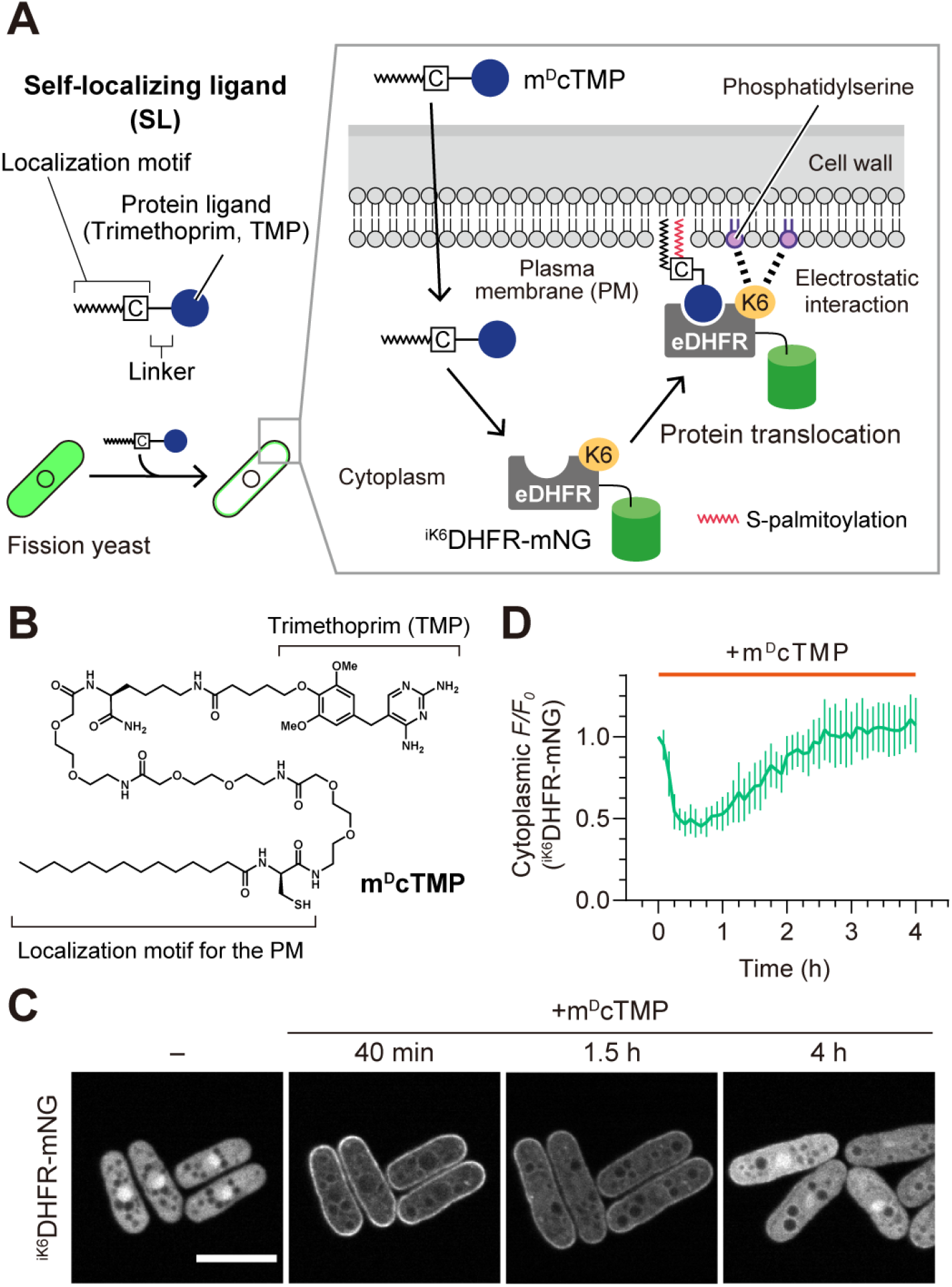
Self-localizing ligand-induced protein translocation (SLIPT) using m^D^cTMP-^iK6^DHFR in *S. pombe*. (A) Schematic illustration of the translocation of an mNeonGreen (mNG)-fused hexalysin (K6 tag)-inserted eDHFR (^iK6^DHFR-mNG) by m^D^cTMP from the cytoplasm to the plasma membrane (PM). The m^D^cTMP and ^iK6^DHFR-mNG complex localizes to the PM through hydrophobic lipid interaction and electrostatic interaction between the K6 tag and phosphatidylserine on the PM. (B) Molecular structure of m^D^cTMP. (C) Confocal fluorescence images of fission yeast cells expressing ^iK6^DHFR-mNG were taken every 5 min. Representative images at the indicated times after addition of m^D^cTMP are shown. Scale bar, 10 μm. (D) Time course of m^D^cTMP-induced ^iK6^DHFR translocation to the PM. Normalized fluorescence intensity in the cytoplasm was plotted as a function of time. Data are presented as mean ± SD (n = 7 cells).

The fission yeast *Schizosaccharomyces pombe* is a widely used eukaryotic model organism. Molecular genetic analysis of *S. pombe* has allowed the identification of genes that regulate cellular processes such as the cell cycle ^23^. However, analysis using gene knockout mutants can only yield results with low temporal resolution, in addition to the difficulty of studying lethal genes. Therefore, it remains a technical challenge to precisely analyze when and how cell cycle-related proteins function in the cell cycle. Although the CID method is expected to provide a way to rapidly perturb intracellular signaling, the FKBP-FRB system in fission yeast has some technical problems, such as the inhibition of sexual development by TOR inhibition with rapamycin ^24–27^. In addition, to use the FKBP-FRB system in fission yeast, the *fkh1* gene must be knocked out due to the high expression levels of endogenous Fkh1 (the FKBP12 homolog) ^24^. For these reasons, there is a need for perturbation tools to control cell signaling with low cytotoxicity and high temporal resolution in fission yeast.

In this study, we applied the SLIPT system to *S. pombe* and successfully demonstrated rapid perturbation of several cell cycle-related proteins. The self-localizing ligands recruited their binding partners to the PM or nucleus in fission yeast. We further optimized the self-localizing ligands for *S. pombe* to maintain recruitment of target molecules to the PM over a long period of time. In addition, we observed the localization changes of dozens of endogenous molecules by knocking in a gene of the binding partner for self-localizing ligands, along with analysis of the associated cell-cycle perturbations and phenotypes. Taken together, we demonstrate for the first time that the SLIPT system serves as a useful chemogenetic tool for rapid perturbation of endogenous molecules in *S. pombe*.

## Results and Discussion

### Introduction of the SLIPT system to fission yeast

In this study, we sought to develop a functional SLIPT system optimized for the fission yeast *S. pombe*. We have previously reported a self-localizing ligand, m^D^cTMP, which is composed of myristoylated D-cysteine (m^D^c) and trimethoprim (TMP) ^17^ (Figure 1B). Within a cell, the cysteine side chain of m^D^cTMP is palmitoylated by palmitoyl acyltransferase at the Golgi apparatus, followed by delivery of the palmitoylated m^D^cTMP to the PM and Golgi ^17,28^ (Figure 1A). *Escherichia coli* dihydrofolate reductase (eDHFR), known to bind specifically to TMP, is redistributed to the PM and Golgi by binding to m^D^cTMP (Figure 1A). Notably, *dfr1*, a dihydrofolate reductase gene in *S. pombe*, is insensitive to TMP ^29^, supporting the orthogonality of this SLIPT system in fission yeast. In addition, we have recently developed ^iK6^DHFR, in which six lysine residues are inserted into the loop region adjacent to the PM of the eDHFR-m^D^cTMP complex ^21^ (Figure 1A). The insertion of the six lysine residues facilitates the selective localization of ^iK6^DHFR to the PM, but not to the Golgi, through electrostatic interactions with the inner leaflet of the PM.

We investigated whether m^D^cTMP could induce relocalization of ^iK6^DHFR from the cytoplasm to the PM in *S. pombe*. The cells expressing ^iK6^DHFR fused to green fluorescent protein mNeonGreen (^iK6^DHFR-mNG) were placed in a microfluidic chamber (Figure S1) and imaged with a spinning disk confocal microscope. ^iK6^DHFR-mNG was diffusely distributed in the cytoplasm and nucleus before the addition of m^D^cTMP (Figure 1C). Addition of 10 μM m^D^cTMP from the microfluidic inlet induced redistribution of ^iK6^DHFR-mNG to the PM within 30 min (Figure 1C and 1D, Movie S1). However, ^iK6^DHFR-mNG did not remain at the PM for a long time, but rather gradually returned to the cytoplasm within approximately 4 h, even in the presence of m^D^cTMP. These results indicate that the m^D^cTMP-eDHFR-based SLIPT system is functional in the fission yeast *S. pombe.* Since eDHFR without an iK6 tag is largely translocated to the endoplasmic reticulum (ER) and Golgi upon addition of m^D^cTMP (Figure S2), the iK6 tag is essential for m^D^cTMP-induced eDHFR translocation to the PM. In addition, this approach would be useful for the transient manipulation of intracellular signaling systems because m^D^cTMP-induced PM translocation of ^iK6^DHFR is transient.

In addition, we tested whether hoeTMP, a self-localizing ligand that recruits DHFR fusion proteins to the nucleus, works in fission yeast. hoeTMP is a small chemical compound consisting of Hoechst 33258 linked to TMP ^15^, and is designed so that the Hoechst 33258 moiety binds selectively to DNA and the DHFR fusion protein is captured in the nucleus via the TMP moiety. We therefore treated fission yeast cells expressing ^iK6^DHFR-mNG with hoeTMP and found that ^iK6^DHFR-mNG localized to the nucleus as expected (Figure S3). These results indicate that hoeTMP functions as a nuclear localizing ligand in fission yeast.

### Improvement of the SLIPT system for fission yeast

We have previously reported that the amide bonds of the original self-localizing ligand, myristoylated glycine-cysteine TMP (mgcTMP), are degraded by proteases within 40 min in mammalian cultured cells, resulting in the spontaneous dissociation of eDHFR from the PM ^17^. m^D^cTMP, an improved version of mgcTMP, is able to maintain plasma membrane localization for more than 24 h in cultured mammalian cells ^17^, whereas in fission yeast, m^D^cTMP-induced PM localization of ^iK6^DHFR was diminished within 1-2 h (Figure 1D). These results suggest that m^D^cTMP is further degraded by protease(s) expressed in fission yeast. To address this issue, we newly synthesized m^D^cTMP_4Me_, in which four amides of m^D^cTMP are *N*-methylated, in order to render it resistant to the protease(s) (Figure 2A), as *N*-methylation of amide bonds is known to increase the stability of compounds ^19^.

**Figure 2.**
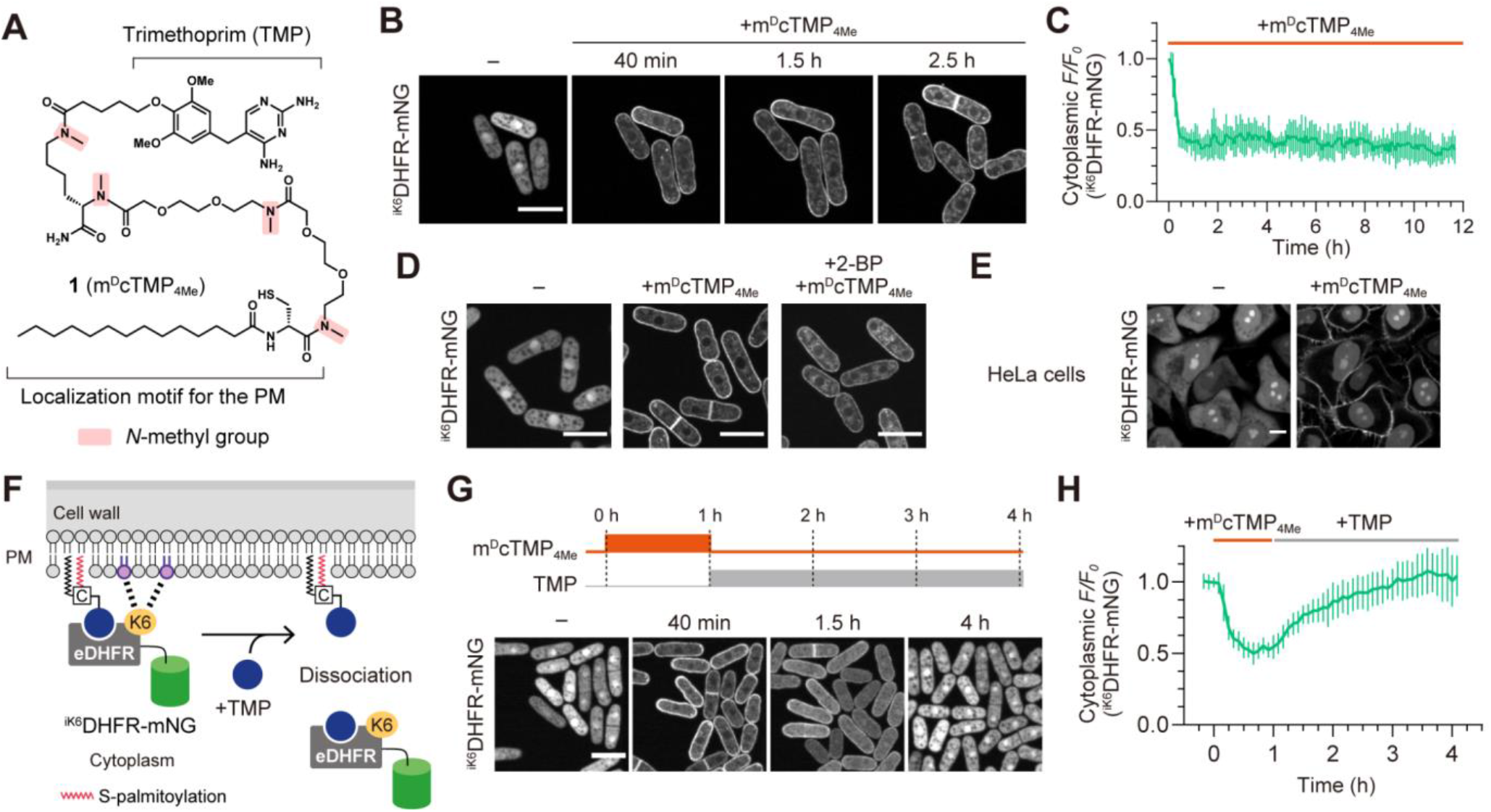
Improvement of the SLIPT system for *S. pombe*. (A) Chemical structure of m^D^cTMP_4Me_. (B and C) Fission yeast cells expressing ^iK6^DHFR-mNG were loaded into a microfluidic device and treated with 10 μM m^D^cTMP_4Me_. (B) Confocal images taken at the indicated time points. Scale bar, 10 μm. (C) Time course of m^D^cTMP_4Me_-induced ^iK6^DHFR translocation to the PM. Normalized fluorescence intensity in the cytoplasm was plotted as a function of time. Data are presented as mean ± SD (n = 7 cells). (D) PM recruitment of ^iK6^DHFR-mNG in the presence of the palmitoylation inhibitor 2-bromopalmitate (2-BP). Confocal images of fission yeast cells expressing ^iK6^DHFR-mNG were taken in the absence of compounds (left) or after treatment with 10 μM m^D^cTMP_4Me_ (middle) or both 10 μM m^D^cTMP_4Me_ and 100 μM 2-BP (right). Scale bar, 10 μm. (E) HeLa cells stably expressing ^iK6^DHFR-mNG were treated with 10 μM m^D^cTMP_4Me_. Confocal fluorescence images were obtained before (left) and 10 min after the addition of m^D^cTMP_4Me_. Scale bar, 10 μm. (F) Schematic of the translocation of PM-localized ^iK6^DHFR-mNG to the cytoplasm by competition with free TMP treatment. (G and H) Fission yeast cells expressing ^iK6^DHFR-mNG were loaded into a microfluidic device and treated with 10 μM m^D^cTMP_4Me_ for 1 h, followed by treatment of 100 μM TMP for 3 h. (G) Confocal fluorescence images of the cells at the indicated time points are shown (lower). Scale bar, 10 μm. (H) Time course of m^D^cTMP_4Me_-induced ^iK6^DHFR translocation to the PM. Normalized fluorescence intensity in the cytoplasm was plotted as a function of time. Data are presented as mean ± SD (n = 10 cells).

Next, we examined whether m^D^cTMP_4Me_ could induce plasma membrane localization of ^iK6^DHFR-mNG in fission yeast and mammalian cells in the manner of m^D^cTMP. ^iK6^DHFR-mNG-expressing fission yeast cells were applied to a microfluidic chamber and m^D^cTMP_4Me_ was added to the cells. ^iK6^DHFR-mNG translocated from the cytoplasm to the PM as expected (Figure 2B, Movie S2). The membrane localization of ^iK6^DHFR-mNG by m^D^cTMP_4Me_ persisted for more than 12 h, though the signal of ^iK6^DHFR-mNG on the endomembrane gradually became prominent approximately 4 h after the addition of m^D^cTMP_4Me_. (Figure 2C, Movie S2). The endomembrane localization of the m^D^cTMP_4Me_-bound ^iK6^DHFR-mNG might be explained by the phosphoatidyl serine-rich surface of the endomembrane in fission yeast ^30^. The membrane localization of ^iK6^DHFR-mNG by m^D^cTMP_4Me_ were reproduced in yeast cells covered with low melting temperature agarose ^31^ (Figure S4). Previous studies have shown that palmitoylation of the cysteine in m^D^cTMP is critical for the plasma membrane selective localization of the m^D^cTMP-^iK6^DHFR-mNG complex ^21^. Therefore, we investigated whether palmitoylation of m^D^cTMP_4Me_ is also important for the plasma membrane localization of ^iK6^DHFR-mNG by m^D^cTMP_4Me_. Fission yeast cells were pretreated with a palmitoylation inhibitor, 2-bromopalmitate (2-BP), followed by the addition of m^D^cTMP_4Me_. ^iK6^DHFR-mNG was localized not only at the plasma membrane but also at endomembranes such as the ER (Figure 2D). This could be due to the fact that m^D^cTMP_4Me_ is localized to the PM and ER only when the myristoyl group of m^D^cTMP_4Me_ remains ^17^. Thus, palmitoylation of m^D^cTMP_4Me_ is essential for PM-selective localization of m^D^cTMP_4Me_-^iK6^DHFR-mNG. Furthermore, treatment of HeLa cells expressing ^iK6^DHFR-mNG with m^D^cTMP_4Me_ resulted in rapid translocation of ^iK6^DHFR-mNG to the PM (Figure 2E), indicating that m^D^cTMP_4Me_ is also functional in cultured cells.

Finally, we attempted to restore ^iK6^DHFR-mNG localization from the PM to the cytoplasm by adding free TMP in fission yeast, as previously accomplished in cultured cells (Figure 2F) ^21^. Fission yeast cells expressing ^iK6^DHFR-mNG were applied to a microfluidic chamber and treated with m^D^cTMP_4Me_ for 1 h, followed by the addition of an excess amount of TMP for 3 h (Figure 2G). We found that ^iK6^DHFR-mNG localized at the plasma membrane gradually returned to the cytoplasm (Figure 2G and 2H). These results demonstrate a potential application of the SLIPT system for transient manipulation of protein localization using free TMP.

### Perturbation of endogenous Cdc2 by using the SLIPT system

To demonstrate the potential of the SLIPT system for manipulating signaling pathways in *S. pombe*, we fused ^iK6^DHFR-mNG to endogenous proteins and attempted to perturb cell signaling by tethering the protein to the plasma membrane with m^D^cTMP_4Me_. We focused on cyclin-dependent kinase (CDK) Cdc2, a key molecule in the cell cycle ^32^. Cell cycle progression is regulated by CDK, which is a nuclear-localized protein and is thought to function primarily in the nucleus ^32^. Therefore, we hypothesized that when endogenous Cdc2 is forced to relocalize to the plasma membrane using m^D^cTMP_4Me_, Cdc2 is excluded from the nucleus, phosphorylation of nuclear substrates is inhibited, and consequently cell cycle progression is blocked.

We established a fission yeast strain in which the ^iK6^DHFR-mNG gene and a selection marker gene were knocked in at the 3’ end of the *cdc2* gene (Cdc2-^iK6^DHFR-mNG) (Figure S5A). The Cdc2-^iK6^DHFR-mNG strain showed no aberrant cell morphology, and the cell cycle duration was nearly 3 h/cycle in the absence of m^D^cTMP_4Me_, similar to that in wild-type cells (Figure 3A and 3B). The cell length at division was approximately 14.5 μm (Figure 3C), which is almost identical to that of the wild-type strain, indicating that the knock-in of the ^iK6^DHFR-mNG gene into the *cdc2* locus had little effect on the cell cycle. Next, m^D^cTMP_4Me_ was added to the Cdc2-^iK6^DHFR-mNG strain, and as expected, almost all the Cdc2-^iK6^DHFR-mNG was exported to from the nucleus and localized to the plasma membrane within 2-4 h (Figure 3D, Movie S3). After the addition of m^D^cTMP_4Me_, the cell length increased in accordance with the prolonged cell cycle duration. Furthermore, several hours after the addition of m^D^cTMP_4Me_, cells with either longer or shorter cell lengths accumulated (Figure 3D and 3E). Notably, longer cells formed multiple septa during cell division (Figure S5B). Finally, the cell cycle was disrupted, resulting in the accumulation of both cells longer and cells shorter than control cells (Figure 3F). As a control experiment, we added m^D^cTMP_4Me_ to a fission yeast strain stably expressing ^iK6^DHFR-mNG and found that the cell cycle duration and cell length at division were not altered (Figure S5C and S5D).

**Figure 3.**
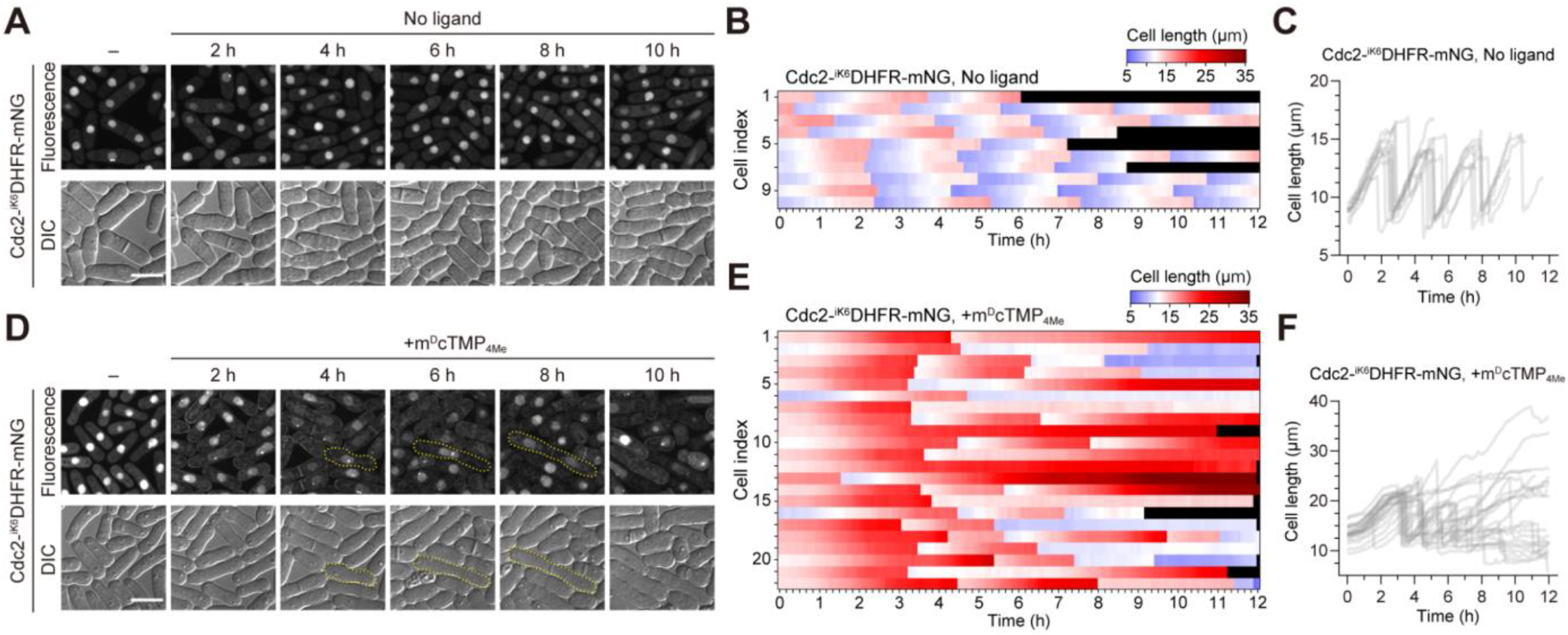
Plasma membrane recruitment of endogenous Cdc2 with the SLIPT system induces cell cycle arrest in *S. pombe*. (A-C) Fission yeast cells expressing ^iK6^DHFR-mNG-fused endogenous Cdc2 were imaged without m^D^cTMP_4Me_ in a microfluidic chamber. (A) Confocal fluorescence images (top) and differential interference contrast (DIC) images (bottom) of fission yeast cells expressing ^iK6^DHFR-mNG-fused endogenous Cdc2 were taken at the indicated time points. (B) A heat map showing the temporal profile of the cell length of fission yeast cells expressing ^iK6^DHFR-mNG-fused endogenous Cdc2 in the absence of m^D^cTMP_4Me_ (n = 10 cells). (C) Cell length in fission yeast cells expressing ^iK6^DHFR-mNG-fused endogenous Cdc2 is plotted as a function of time (n = 10 cells). (D-F) Fission yeast cells expressing ^iK6^DHFR-mNG-fused endogenous Cdc2 were imaged with m^D^cTMP_4Me_ in a microfluidic chamber. (D) Confocal fluorescence images (top) and DIC images (bottom) of fission yeast cells expressing ^iK6^DHFR-mNG-fused endogenous Cdc2 were taken at the indicated time points after m^D^cTMP_4Me_ treatment. (E) A heat map showing the temporal profile of the cell length of fission yeast cells expressing ^iK6^DHFR-mNG-fused endogenous Cdc2 in the presence of m^D^cTMP_4Me_ (n = 22 cells). (F) Cell length in fission yeast cells expressing ^iK6^DHFR-mNG-fused endogenous Cdc2 is plotted as a function of time after m^D^cTMP_4Me_ treatment (n = 22 cells).

When endogenous Cdc2 is forced to be recruited to the plasma membrane using the SLIPT system, fission yeast cells eventually exhibit two phenotypes: one group of cells that continue to elongate and another group of cells that maintain a short cell length (Figure 3F). In addition, the recruitment of Cdc2 to the plasma membrane often caused the formation of multiple septations in longer cells (Figure S5B). This may have been because the SLIPT system dysregulated the Septation Initiation Network (SIN) mechanism by recruiting Cdc2 proteins ^33^. Of note, the kinetics of plasma membrane translocation of Cdc2-^iK6^DHFR-mNG by m^D^cTMP_4Me_ was slower (2-4 h) than that of ^iK6^DHFR-mNG (15-30 min) (Figures 2C and 3D). This could be due to the fact that Cdc2-^iK6^DHFR-mNG must first be translocated from the nucleus to the cytoplasm, before it can be translocated to the plasma membrane by m^D^cTMP_4Me_.

### Perturbation of cell cycle-related proteins with the SLIPT system in fission yeast

Finally, we used the SLIPT system to perturb cell cycle-associated proteins and examine their phenotypes. The genes we targeted for SLIPT are listed in Table 1. We established fission yeast strains in which the *^iK6^DHFR-mNG* gene was knocked in at the 3’ end of each target gene. In the absence of m^D^cTMP_4Me_, no noticeable phenotypes were observed for any of the strains, with the exception that the Sfi1-^iK6^DHFR-mNG strain exhibited a slower growth rate than the WT strain. These cell lines were treated with m^D^cTMP_4Me_ (10 μM), incubated for 12 h with shaking, and observed under a fluorescence microscope (Figure 4A). The phenotypes are summarized in Table 1.

**Figure 4.**
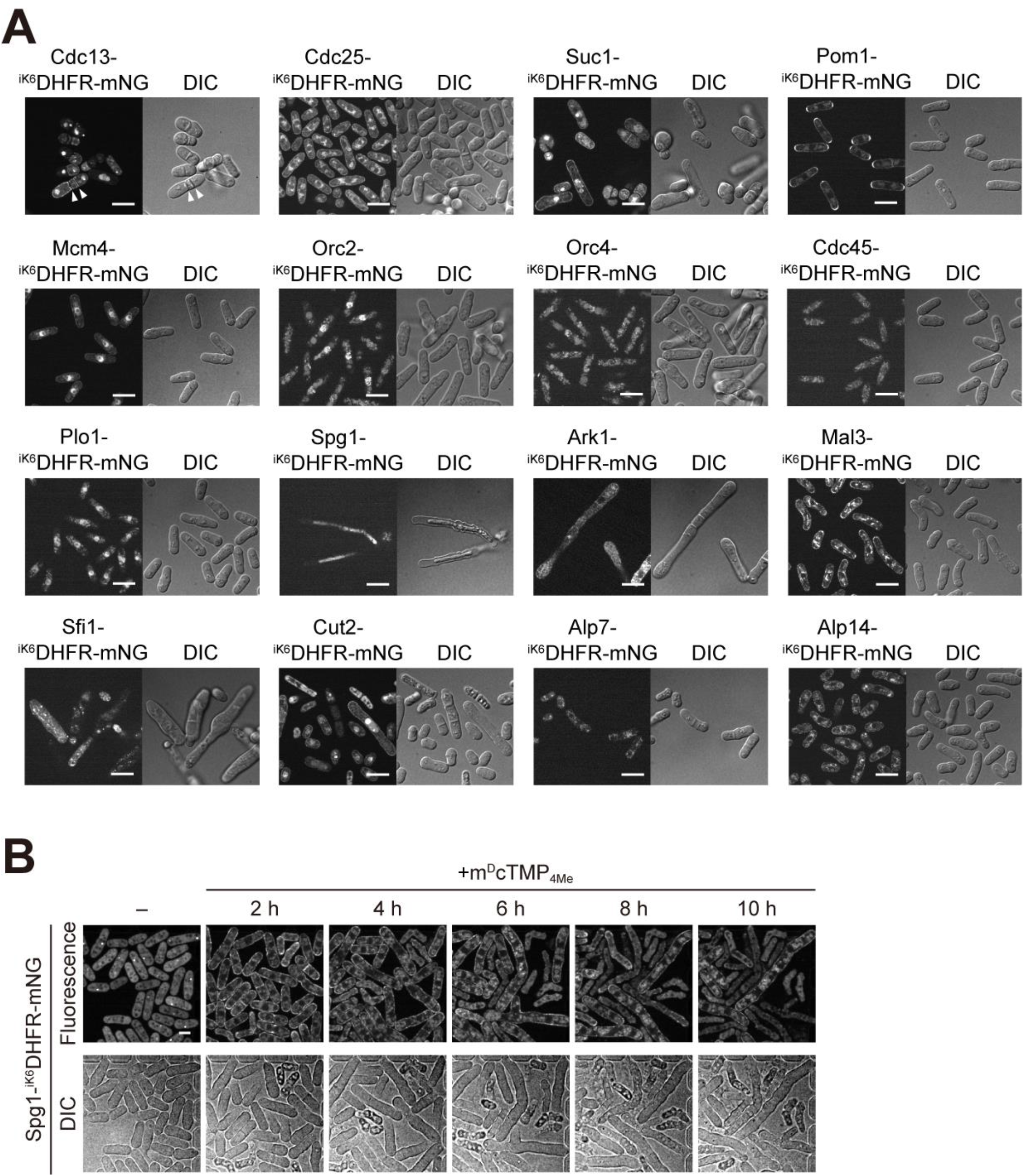
Perturbation of cell cycle-related proteins using the SLIPT system. (A) Confocal (left) and DIC (right) images of ^iK6^DHFR-mNG knock-in strains 12 h after treatment with 10 μM m^D^cTMP_4Me_. (B) Fission yeast cells expressing ^iK6^DHFR-mNG-fused endogenous Spg1 were time-lapse imaged with m^D^cTMP_4Me_ in a microfluidic chamber. Confocal fluorescence images (top) and DIC images (bottom) of fission yeast cells expressing ^iK6^DHFR-mNG-fused endogenous Spg1 were taken at the indicated time points.

**Table 1.** Phenotypes induced by the SLIPT system.

| Target gene | Product | SLIPT phenotype |
| --- | --- | --- |
| *cdc13* | B-type cyclin | Shorter cell length, multi-septum |
| *cdc25* | Phosphatase, M-phase inducer | No obvious phenotype |
| *suc1* | CDK regulatory subunit | Shorter cell length, multi-septum |
| *mcm4* | MCM complex subunit | No obvious phenotype |
| *orc2* | Origin recognition complex subunit | Slightly longer cell length |
| *orc4* | Origin recognition complex subunit | Slightly longer cell length |
| *cdc45* | DNA replication pre-initiation complex subunit | No obvious phenotype |
| *plo1* | Polo kinase | No obvious phenotype |
| *pom1* | DYRK family cell polarity protein kinase | No obvious phenotype |
| *spg1* | Septum Promoting GTPase | Longer cell length, cell death |
| *ark1* | Aurora-B kinase | Longer cell length |
| *mal3* | Microtubule plus-end binding protein, EB1 family | Banana shape |
| *sfi1* | Spindle pole body half bridge protein | Morphological defect |
| *cut2* | Sister chromatid separation inhibitor, securin | Shorter and longer cell length, multi septum |
| *alp7* | TACC protein | No obvious phenotype |
| *alp14* | TOG/XMAP215 microtubule plus-end tracking polymerase | No obvious phenotype |

Upon m^D^cTMP_4Me_ treatment, Cdc13-^iK6^DHFR-mNG and Suc1-^iK6^DHFR-mNG cells showed shorter cell lengths than WT cells, and multiple septations were observed in some cells (Figure 4A, white arrowheads). These results were close to the phenotype of m^D^cTMP_4Me_-treated Cdc2-^iK6^DHFR-mNG cells, suggesting that endogenous Cdc2 was also recruited to the membrane and caused the dysregulation of the SIN mechanism. In contrast, no remarkable phenotype was observed by the SLIPT system for the other Cdc2 regulators, *i.e.*, Cdc25 and Pom1 (Figure 4A). Mcm4, Orc2, Orc4, and Cdc45, which are associated with S phase progression, showed little phenotype by the SLIPT system. These results might have been due to insufficient membrane recruitment of these factors as shown in the mNG fluorescence images (Figure 4A), which implyied that these factors are tightly bound to histones and/or DNA and therefore are not affected by the SLIPT system. Finally, genes related to mitosis and cytokinesis were perturbed by SLIPT. m^D^cTMP_4Me_ treatment killed Spg1-^iK6^DHFR-mNG cells with elongated cell length (Figure 4A). To confirm when and how the Spg1-^iK6^DHFR-mNG cells died, we performed time-lapse imaging. We found that m^D^cTMP_4Me_ treatment rapidly induced membrane translocation of Spg1-^iK6^DHFR-mNG from the spindle pole body (SPB) and cytoplasm, resulting in sustained cell length elongation, multiple nuclear divisions without cytokinesis, and gradual cell death (Figure 4B, Movie S4). This result is consistent with the phenotype of the Spg1ts mutant ^34,35^. Upon m^D^cTMP_4Me_ treatment, Ark1-^iK6^DHFR-mNG and Cut2-^iK6^DHFR-mNG cells exhibited a mixture of long and short cells, whereas Mal3-^iK6^DHFR-mNG and Sfi1-^iK6^DHFR-mNG cells showed a banana shape and morphological defects, respectively (Figure 4A). On the other hand, no significant phenotype was observed for the perturbation of Pol1, Alp7, or Alp14 (Figure 4A), due to there being only partial or no membrane translocation of these proteins by the SLIPT system.

In this study, we succeeded in altering the localization of the ^iK6^DHFR fusion protein by a self-localizing ligand in fission yeast cells, and demonstrated that this SLIPT system enables perturbation of the localization and functions of several molecules associated with the cell cycle. Temperature-sensitive mutants, which have been widely used in genetic studies of fission yeast, take a long time to show a phenotype after a change in temperature, whereas the SLIPT system could rapidly perturb the localization and function of molecules. In addition, the SLIPT system does not require excitation light, allowing simultaneous fluorescence imaging and multiple perturbations in combination with optogenetic tools. Compared to other CID systems, such as the FKBP-FRB system, the SLIPT system in fission yeast has several advantages. First, the FKBP-FRB system requires fusion of FKBP and FRB to the target and localized molecules, respectively, whereas the SLIPT system only requires fusion of ^iK6^DHFR to the target molecule. Second, unlike the FKBP-FRB system, the SLIPT system can reversibly manipulate localization ^36^.

Several technical challenges remain for the fission yeast SLIPT system. First, the ^iK6^DHFR fusion molecule is completely localized to the plasma membrane by m^D^cTMP in approximately 5 min in cultured mammalian cells ^21^, but it took approximately 15-30 min for the ^iK6^DHFR-mNG to translocate to the plasma membrane in fission yeast (Figure 1D and 2C). This result may have been be due to the lower permeability of localized ligands through the cell wall and/or plasma membrane and the higher activity of multidrug transporters in fission yeast ^37^. This issue could be addressed by synthesizing ligands with higher cell wall and/or plasma membrane permeability. Second, self-localizing ligands for intracellular organelles other than the plasma membrane and nucleus have not been tested in fission yeast. In this study, we developed and tested a localizing ligand that targeted the DHFR fusion protein to the plasma membrane and nucleus in fission yeast. In cultured cells, self-localizing ligands for Golgi apparatus, endoplasmic reticulum, and microtubules have been reported ^15,18,19^. Reuse of these ligands will make them more useful chemical and genetic tools in fission yeast. Third, we found that SLIPT-induced localization changes could not be observed for the factors that were tightly bound to intracellular organelles and components, such as Mcm4 (Figure 4A). This may be because the affinity between the self-localizing ligand and the DHFR-fused endogenous molecule was not sufficient to dissociate the endogenous molecule from the intracellular organelle. Indeed, a previous report showed that the localization of endogenous Mcm4 was altered in a CID system using FKBP-FRB ^24^, which induced irreversible heterodimerization.

Finally, we would like to suggest some future directions for SLIPT systems using fission yeast. First, more rapid and localized perturbation could be achieved by adding caged moieties to self-localizing ligands. m^D^cTMP^NVOC^, a photoactivatable self-localizing ligand that can uncage m^D^cTMP with UV, has already been reported ^20,38^. The introduction of four *N*-methylations into m^D^cTMP^NVOC^ will allow the molecule to be perturbed with high temporal and spatial resolution by local light illumination in fission yeast. Secondly, genome-wide perturbation and its phenotypic analysis could be useful direction for SLIPT. A library of fission yeast strains with the *GFP* gene knocked in at the 3’ end of the endogenous gene is available ^39^, and thus the expression of DHFR-fused GFP nanobodies ^40^ with self-localizing ligands will enable efficient genome-wide manipulation of endogenous molecules. The fission yeast SLIPT system will provide new opportunities for understanding various biological phenomena.

## Methods

### Plasmid construction

All plasmids used in this study are listed in Supplemental Table S1 with Benchling links that provides the sequences and plasmid maps. pNATZA11 [National Bio-Resource Project (NBRP) plasmid FYP2875, provided by Dr. Watanabe (The University of Tokyo) and pFA6a-mNG-kan [NBRP plasmid FYP4997] ^41^ were used as vector backbones to construct plasmids for the establishment of stable cell lines of the fission yeast *S. pombe*. pCSIIpuro-mNG-eDHFR(69K6) ^21^ was used as a PCR template and lentiviral vector for the establishment of a stable mammalian cell line. All plasmids were generated using standard cloning procedures and the NEBuilder HiFi DNA assembly system (New England Biolabs).

### Culture and transformation of fission yeast *S. pombe*

A wild-type fission yeast (L972) was obtained from the NBRP (FY7507, provided by Dr. Shimoda, Osaka City University). The growth medium and other techniques for fission yeast were based on the protocol described previously ^42^, unless otherwise noted. The transformation protocol was modified from the previous report ^43^. In brief, L972 cells were precultured in 2 mL SD medium [0.67% (w/v) bacto yeast nitrogen base without amino acids (Becton, Dickinson and Co.), 0.5% (w/v) D(+)-glucose] at 32°C. After centrifugation at 860 × g, 1 μL of sheared salmon sperm DNA (BioDynamics Lab. Inc.), 10-100 ng of linear DNA encoding the left/right homology arms, and the protein expression cassettes were added. The protein expression cassettes include the cDNA of the protein of interest (POI) and a drug-resistance gene. After resuspending the cells in a mixture of 50 μL LiOAc/TE buffer (100 mM LiOAc, 10 mM Tris-HCl, 1 mM EDTA, pH 7.6) containing 30% (v/v) glycerol and 145 μL LiOAc/TE buffer containing 50% (w/v) polyethylene glycol 4000 (FUJIFILM Wako Pure Chemical Corp.), the suspension was incubated at 42°C for 10 min. After centrifugation at 860 × g, the cells were resuspended in 20 μL distilled water and plated on the YEA agar plate [0.5% (w/v) yeast extract (Nacalai Tesque), 3% (w/v) D(+)-glucose, 1.67 mM adenine (FUJIFILM Wako Pure Chemical Corp.), 2% (w/v) agar (Nacalai Tesque)] and incubated at 32°C for at least 12 h. The cells were then transferred to the new YEA agar plate containing antibiotics [100 μg/mL G418 (InvivoGen) or 100 μg/mL nourseothricin (Gold Biotechnology)] and incubated at 32°C for at least 3 days to select the transformants. The expected protein expression in the obtained single clones was confirmed by fluorescence imaging. All cell lines used in this study are listed in Supplemental Table S2.

### Live cell imaging

Confocal fluorescence imaging was performed on an IX83 equipped with a UPALXAPO60XO/1.42 NA oil objective lens (Olympus), a UPALXAPO100XO/1.45 NA oil objective lens (Olympus), a Z-drift compensator system (IX3-ZDC2, Olympus), a sCMOS camera (Orca-Fusion BT; Hamamatsu Photonics), and a spinning disk confocal unit (CSU-W1; Yokogawa Electric Corp.). For fission yeast time-lapse imaging, a CellASIC ONIX2 Microfluidic System (CAX2-S0000, Millipore) and a CellASIC ONIX2 Manifold XT (CAX2-MXT20, Millipore) were used. The microscope was controlled by MetaMorph software (Molecular Devices). A 488 nm laser was used for excitation of mNeonGreen. An excitation dichroic mirror (DM405/488/561/640) and emission filter (525/50 for mNeonGreen) were used for confocal fluorescence imaging (Yokogawa Electric Corp.). Fluorescence images were analyzed using Fiji/ImageJ ^44^.

### SLIPT assay in fission yeast *S. pombe*

Established cell lines were precultured in YEA medium [0.5%(w/v) yeast extract (Nacalai Tesque), 3%(w/v) D(+)-glucose, 1.67 mM adenine] at 32°C. For time-lapse imaging, the precultured cells were diluted with fresh YEA medium and loaded into a PDMS-based microfluidic chamber (Y04C-02 or Y04T-04, Millipore). Cells were monitored by time-lapse imaging before and after starting continuous loading with m^D^cTMP (10 μM), m^D^cTMP_4Me_ (10 μM), or TMP (100 μM) at 68.9 kPa.

For perturbation of cell cycle-related proteins, established cell lines were precultured in 2 mL YEA medium at 32°C, followed by the addition of m^D^cTMP_4Me_ (10 μM) and incubation at 32°C for 12 h. For live cell imaging, the cells were seeded on 35 mm glass-base dishes (D11531H, Matsunami Glass). For the SLIPT assay using m^D^cTMP, fission yeast cells expressing eDHFR-mNG were precultured in 2 mL YEA medium, followed by the addition of m^D^cTMP (10 μM) and incubation at 32°C for 30 min. For live cell imaging, the cells were seeded on 35 mm glass-base dishes (Matsunami Glass). For the SLIPT assay using hoeTMP, fission yeast cells expressing ^iK6^DHFR-mNG were precultured in 2 mL YEA medium, followed by the addition of hoeTMP (10 μM) and incubation at 32°C for 60 min. For live cell imaging, the cells were seeded on 35 mm glass-base dishes (Matsunami Glass).

### Palmitoylation assay

Fission yeast cells expressing ^iK6^DHFR-mNG were precultured in 2 mL YEA medium at 32°C, followed by the addition of m^D^cTMP_4Me_ (10 μM) and incubation at 32°C for 40 min. For palmitoylation inhibition, cells were treated with 100 μM 2-bromopalmitate (FUJIFILM Wako Pure Chemical Corp.) and additionally cultured at 32°C for 40 min. For live cell imaging, the cells were seeded on 35 mm glass-base dishes (Matsunami Glass).

### SLIPT assay in fission yeast *S. pombe* on an agarose pad

Fission yeast cells were preincubated in YEA medium at 32°C and plated at 4.0 × 10^5^ cells in the central area of 35 mm glass-base dishes coated with poly-L-lysine (Matsunami Glass). After incubation for 15 min at room temperature, the supernatant was removed, and 100 μL of YEA medium containing 1% (w/v) microwave-heated low-melting point agarose (SeaPlaque agarose, 50101; Lonza) was gently added to the central area. After resting for 15 minutes at room temperature, the cells were observed by time-lapse imaging, and 10 μM m^D^cTMP_4Me_ in 2 mL YEA was added to the dish.

### Mammalian cell culture and establishment of stable cell lines

HeLa cells were obtained from the Cell Resource Center for Biomedical Research, Institute of Development, Aging and Cancer, Tohoku University. Lenti-X-293T cells were purchased from Clontech Laboratories. All cells were maintained in Dulbecco’s modified Eagle’s medium (DMEM) (Nacalai Tesque) supplemented with 10% heat-inactivated fetal bovine serum (FBS) (Sigma), 100 U/mL penicillin, and 100 μg/mL streptomycin (Nacalai Tesque) at 37°C under a humidified 5% CO_2_ atmosphere. For live cell imaging, cells were seeded on 35 mm glass-base dishes (Iwaki) coated with collagen type I-C (Nitta Gelatin).

For lentiviral production, Lenti-X-293T cells were cotransfected with the pCSIIpuro-mNG-eDHFR(69K6), psPAX2 (Addgene plasmid #12260, a gift from D. Trono), and pCMV-VSV-G-RSV-Rev (RIKEN BioResource Research Center plasmid RDB04393, provided by Dr. Miyoshi, Keio University) plasmids ^45^ using a polyethlenimine “Max” (Polysciences Inc.). Virus-containing media were collected 48 h after transfection, filtered, and used to infect target HeLa cells with 8 μg/mL polybrene (Nacalai Tesque). Infected HeLa cells were selected with 0.5 μg/mL puromycin (InvivoGen) for at least 7 days. Bulk populations of selected cells were used.

### SLIPT assay in mammalian cells

HeLa cells stably expressing mNG-^iK6^DHFR were plated at 1.5 × 10^5^ cells in collagen-coated 35 mm glass-base dishes and cultured at 37°C in 5% CO_2_ for 24 h. The medium was changed to serum-free DMEM (Gibco) supplemented with 100 U/mL penicillin and 100 μg/mL streptomycin, and the cells were observed by time-lapse imaging before and after the addition of m^D^cTMP_4Me_ (10 μM).

## Author Information

### Corresponding Authors

Kazuhiro Aoki - Quantitative Biology Research Group, Exploratory Research Center on Life and Living Systems (ExCELLS), National Institutes of Natural Sciences, 5-1 Higashiyama, Myodaiji-cho, Okazaki, Aichi 444-8787, Japan; Division of Quantitative Biology, National Institute for Basic Biology, National Institutes of Natural Sciences, 5-1 Higashiyama, Myodaiji-cho, Okazaki, Aichi 444-8787, Japan; Department of Basic Biology, School of Life Science, SOKENDAI (The Graduate University for Advanced Studies), 5-1 Higashiyama, Myodaiji-cho, Okazaki, Aichi 444-8787, Japan.

Yuhei Goto - Quantitative Biology Research Group, Exploratory Research Center on Life and Living Systems (ExCELLS), National Institutes of Natural Sciences, 5-1 Higashiyama, Myodaiji-cho, Okazaki, Aichi 444-8787, Japan; Division of Quantitative Biology, National Institute for Basic Biology, National Institutes of Natural Sciences, 5-1 Higashiyama, Myodaiji-cho, Okazaki, Aichi 444-8787, Japan; Department of Basic Biology, School of Life Science, SOKENDAI (The Graduate University for Advanced Studies), 5-1 Higashiyama, Myodaiji-cho, Okazaki, Aichi 444-8787, Japan.

### Authors

Akinobu Nakamura - Quantitative Biology Research Group, Exploratory Research Center on Life and Living Systems (ExCELLS), National Institutes of Natural Sciences, 5-1 Higashiyama, Myodaiji-cho, Okazaki, Aichi 444-8787, Japan.

Hironori Sugiyama - Quantitative Biology Research Group, Exploratory Research Center on Life and Living Systems (ExCELLS), National Institutes of Natural Sciences, 5-1 Higashiyama, Myodaiji-cho, Okazaki, Aichi 444-8787, Japan.

Shina Tsukiji - Department of Life Science and Applied Chemistry, Nagoya Institute of Technology, Gokiso-cho, Showa-ku, Nagoya 466-8555, Japan; Department of Nanopharmaceutical Sciences, Nagoya Institute of Technology, Gokiso-cho, Showa-ku, Nagoya 466-8555, Japan.

### Author Contributions

A.N.,Y.G., and K.A conceived the study. A.N., and S.T. synthesized m^D^cTMP, m^D^cTMP_4Me_, and hoeTMP. A.N., H.S., and Y.G. performed the plasmid construction, transformation and establishment of fission yeast strains, time-lapse imaging, and data analysis. A.N. graphed and analyzed the data. A.N., Y.G., S.T., and K.A. drafted the manuscript. All authors approved the manuscript.

### Conflict of Interest

The authors declare no competing financial interest.

## Acknowledgments

We thank Dr. Keiko Kuwata (Nagoya University) for HRMS measurement of m^D^cTMP_4Me_. We thank all members of the Aoki laboratory for their helpful discussions and assistance. Some fission yeast strains were provided by the National Bio-Resource Project (NBRP), Japan. A.N. was supported by JSPS Research Fellowship for Young Scientists (no. JP19J01341). K.A. was supported by a CREST, JST grant (JPMJCR1654) and JSPS KAKENHI grants (nos. JP19H05798 and JP22H02625). Y.G. was supported by a JST, ACT-X Grant Number JPMJAX22B8, JSPS KAKENHI grants (nos. JP19K16050 and JP22K15110), and a Sumitomo Research grant. H.S. was supported by JSPS KAKENHI Grants (nos. JP21J01354 and JP22K15115) and the Dr. Yoshifumi Jigami Memorial Fund, The Society of Yeast Scientists. S.T. was supported by JSPS KAKENHI grants (nos. JP18H04546 and JP20H04706 for the innovative area “Chemistry for Multimolecular Crowding Biosystems”, and JP21H05226 Bottom-up Biotech).

## Notes

### Competing Interest Statement

The authors have declared no competing interest.

### Summary of Updates

Figure S4 was moved to the Supplementary Methods section

