## Supporting Information for "Chemogenetic manipulation of endogenous proteins in fission yeast using a self-localizing ligand-induced protein translocation (SLIPT) system"

### Supplementary Figures

### Figure S1

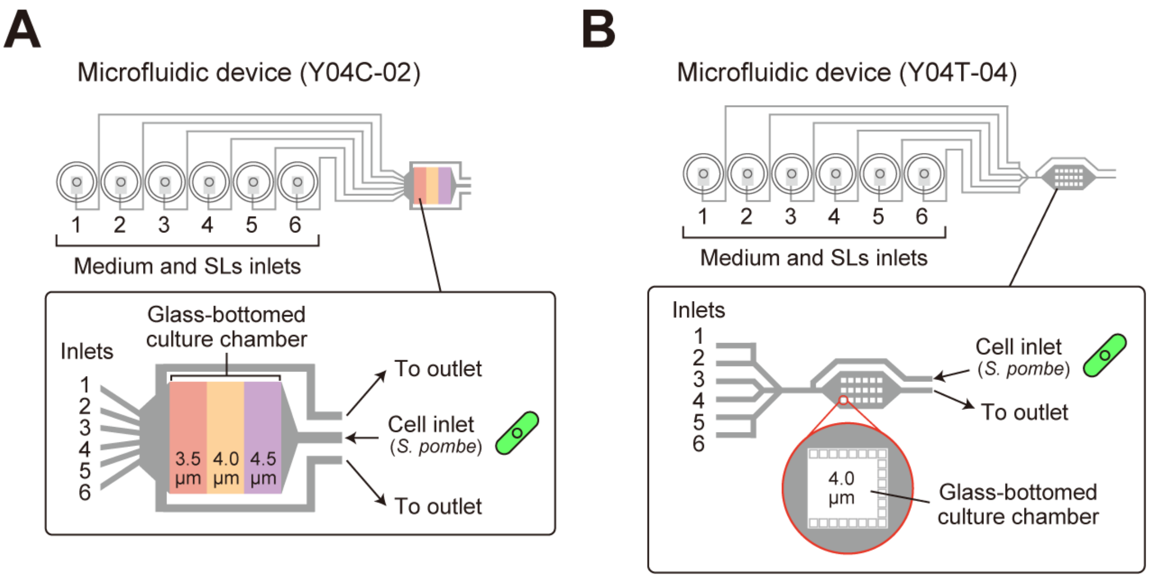


**Figure S1. Microfluidic devices used in this study.**

(A and B) Schematic representation of the microfluidic devices used in this study.

(A) In the microfluidic device (Y04C-02), fission yeast cells are loaded from the cell inlet and sandwiched between the chamber and the glass bottom in the middle region. Culture medium is supplied from inlets 1 to 6.

(B) In the microfluidic device (Y04T-04), fission yeast cells are loaded from the cell inlet, and trapped in the glass-bottomed culture chamber. Culture medium is supplied from inlets 1 to 6.

#### Figure S2

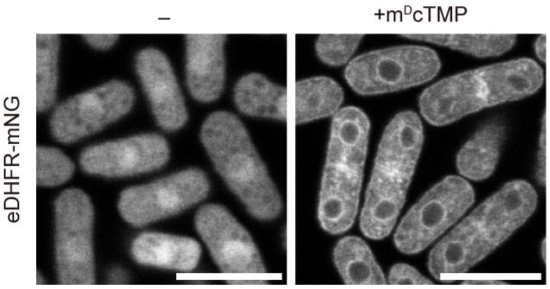


**Figure S2. Endomembrane translocation of the eDHFR fusion protein by m$^{D}$cTMP.**

Confocal fluorescence images of fission yeast cells expressing eDHFR-mNG before (left) and after 30 min (right) of treatment with m$^{D}$cTMP (10 μM). Scale bars, 10 μm.

### Figure S3

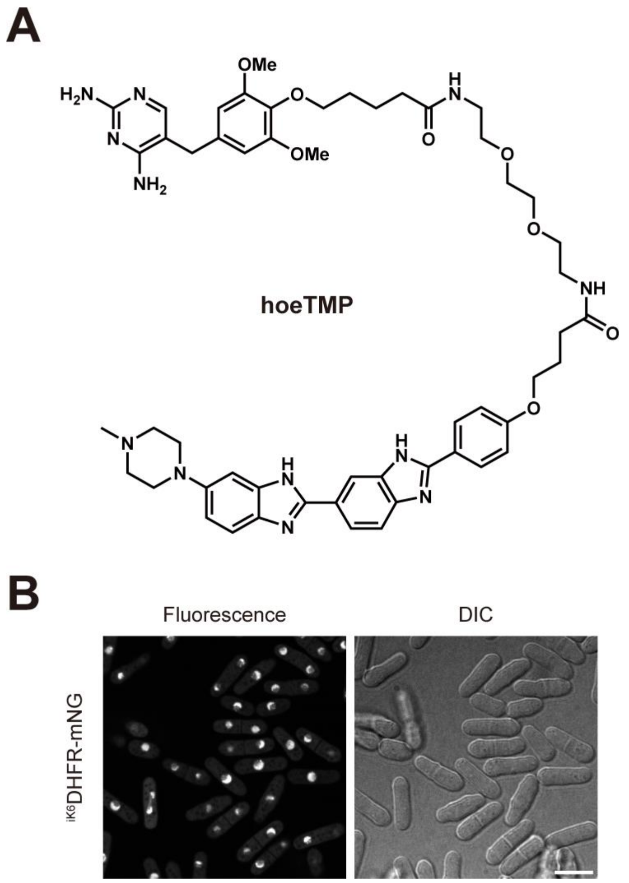


**Figure S3. Nuclear translocation of the DHFR fusion protein by hoeTMP.**

(A) Chemical structure of hoeTMP.

(B) Confocal fluorescence images (left) and DIC image (right) of fission yeast cells expressing $^{iK6}$DHFR-mNG after 60 min of treatment with hoeTMP (10 μM). Scale bar, 10 μm.

### Figure S4

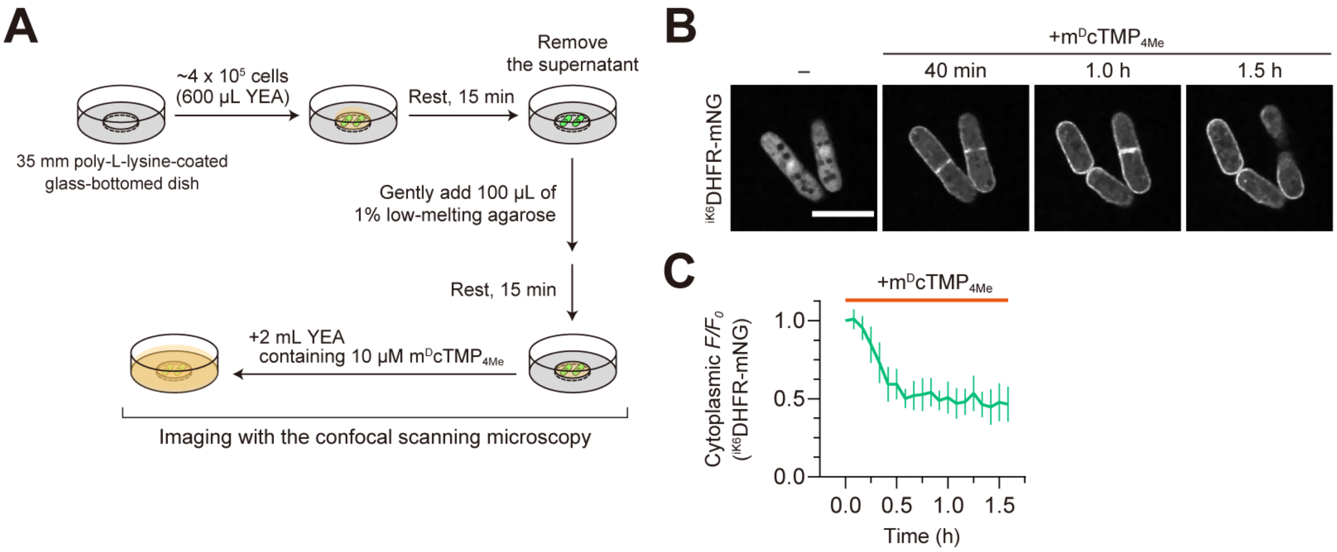


**Figure S4. Time-lapse imaging of fission yeast using an agar pad.**

(A) The procedure for time-lapse imaging of fission yeast using an agar pad is shown.

(B) Confocal fluorescence images of fission yeast cells expressing $^{iK6}$DHFR-mNG were taken every 5 min. The representative images are shown at the indicated time points after the addition of 10 µM m$^{D}$cTMP$_{4Me}$. Scale bar, 10 µm.

(C) Time course of m$^{D}$cTMP$_{4Me}$-induced $^{iK6}$DHFR-mNG translocation to the PM. Normalized fluorescence intensity in the cytoplasm was plotted as a function of time. Data are presented as mean ± SD (n = 6 cells).

**Figure S5**

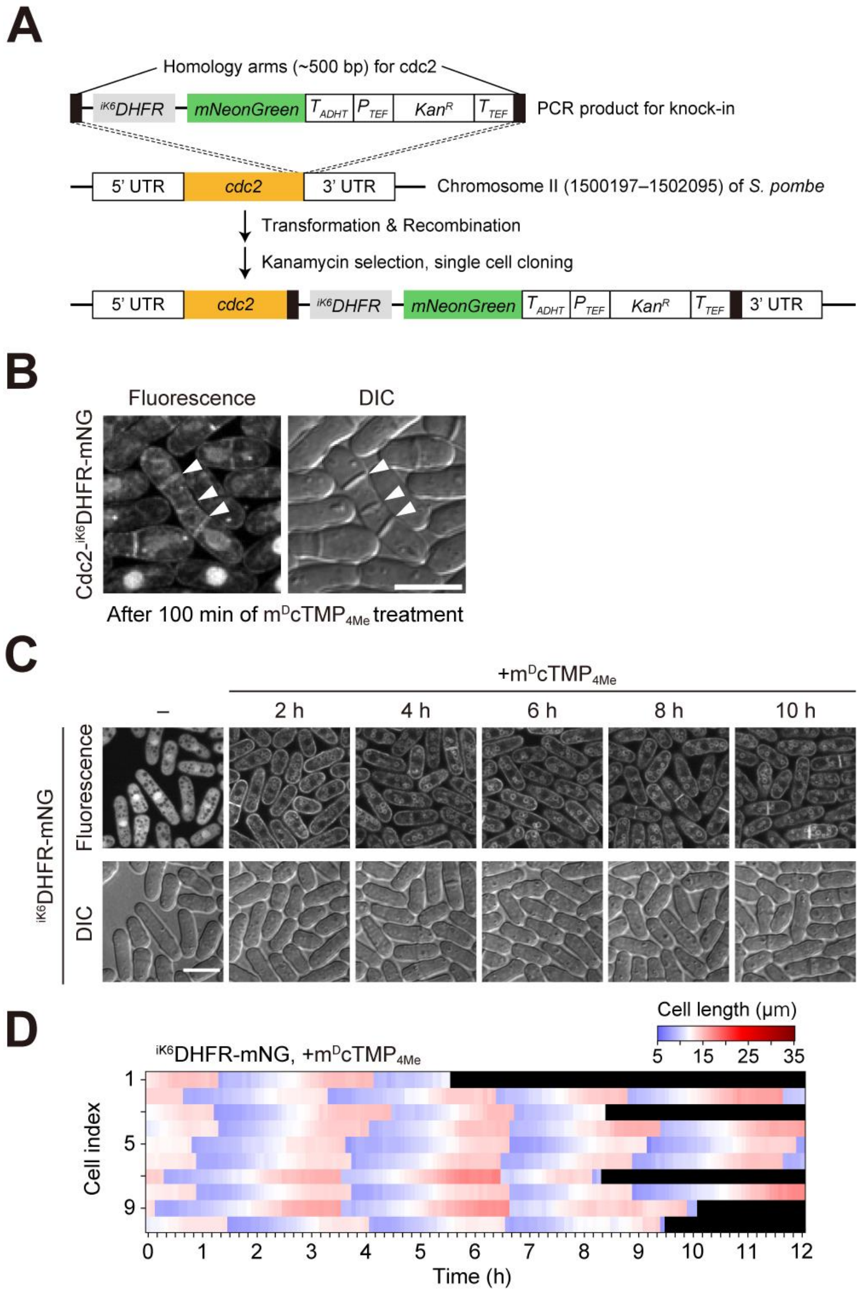


**Figure S5. Endogenous tagging of $^{iK6}$DHFR to the *cdc2* gene.**

(A) Schematic representation of endogenous tagging of $^{iK6}$DHFR to the *cdc2* gene.

(B) Multiple septa formation in fission yeast cells expressing Cdc2-$^{iK6}$DHFR-mNG after 100 min of treatment with m$^{D}$cTMP$_{4Me}$ (10 μM). White arrowheads indicate the septum positions. Scale bar, 10 μm.

(C) Confocal fluorescence images (top) and DIC images (bottom) of fission yeast cells expressing $^{iK6}$DHFR-mNG were taken every 5 min. The representative images are shown at the indicated time after the addition of 10 μM m$^{D}$cTMP$_{4Me}$. Scale bar, 10 μm.

(D) A heat map showing the temporal profile of the cell length of fission yeast cells expressing $^{iK6}$DHFR-mNG in the presence of m$^{D}$cTMP$_{4Me}$ (n = 10 cells).

#### Supplementary Movies

**Movie S1. m$^{D}$cTMP-induced transient membrane translocation of $^{iK6}$DHFR-mNG in fission yeast.** Fission yeast cells expressing $^{iK6}$DHFR-mNG were added to 10 μM m$^{D}$cTMP, and time-lapse imaged. Scale bar, 10 μm.

**Movie S2. m$^{D}$cTMP$_{4Me}$-induced sustained membrane translocation of $^{iK6}$DHFR-mNG in fission yeast.** Fission yeast cells expressing $^{iK6}$DHFR-mNG were added to 10 μM m$^{D}$cTMP$_{4Me}$, and time-lapse imaged. Scale bar, 10 μm.

**Movie S3. m$^{D}$cTMP$_{4Me}$-induced translocation of endogenous Cdc2 for the plasma membrane.** Fission yeast cells expressing Cdc2-$^{iK6}$DHFR-mNG were added to 10 μM m$^{D}$cTMP$_{4Me}$, and time-lapse imaged. Scale bar, 10 μm.

**Movie S4. m$^{D}$cTMP$_{4Me}$-induced translocation of endogenous Spg1 for the plasma membrane.** Fission yeast cells expressing Spg1-$^{iK6}$DHFR-mNG were added to 10 μM m$^{D}$cTMP$_{4Me}$, and time-lapse imaged. Scale bar, 10 μm.

### Supplementary Tables

#### Table S1. Plasmids used in this study.

| **Plasmid name** | **Description** | **Source** | **Benchling Link** |
|---|---|---|---|
| pNATZA11 | pNATZA | NBRP plasmid, FYP2875 | https://benchling.com/s/seq-4c60IcMVvdr5idH0B0Ns |
| pNATZA11-eDHFR-HA-mNeonGreen | pNATZA-eDHFR-mNG | This study | https://benchling.com/s/seq-uU2ogqoRlEgvJJSxRjl3?m=slm-3k71VBk2nVCGv0dkB8j6 |
| pNATZA11-eDHFR(69K6)-HA-mNeonGreen | pNATZA-eDHFR(69K6)-mNG | This study | https://benchling.com/s/seq-aCh5TVHtQ3pY6ejlAohO |
| pFA6a-iRFP-kan | pFA6a-iRFP-kan | NBRP plasmid, FYP4993 | https://benchling.com/s/seq-bkYIYdFO0LlSWLRgqKf2 |
| pFA6a-eDHFR(69K6)-HA-mNeonGreen-kan | pFA6a-eDHFR(69K6)-mNG-kan | This study | https://benchling.com/s/seq-OVK5sJAXdYKzbqM0hKvD |

*kan: kanamycin resistance gene (kanMX6).

*mNeonGreen: the codon-optimized mNeonGreen for *Schizosaccharomyces pombe*.

*eDHFR(69K6): $^{iK6}$DHFR.

**Table S2. Fission yeast strains used in this study.**

| Strain name | Genotype | Figure | Source |
|---|---|---|---|
| L972 | h- | | NBRP, FY7507 |
| AN002 | h- z::Padh11-eDHFR-HA-mNeonGreen<<nat | Fig. S2 | This study, L972 |
| AN054 | h- z::Padh11-eDHFR(69K6)-HA-mNeonGreen<<nat | Fig. 1, Fig.2, Fig.S3, Fig.S4, Fig.S5C | This study, L972 |
| AN029 | h- cdc2-eDHFR(69K6)-HA-mNeonGreen<<kan | Fig. 2, Fig.3, Fig.S5B | This study, L972 |
| AN030 | h- cdc13-eDHFR(69K6)-HA-mNeonGreen<<kan | Fig. 4 | This study, L972 |
| AN031 | h- cdc25-eDHFR(69K6)-HA-mNeonGreen<<kan | Fig. 4 | This study, L972 |
| AN032 | h- mcm4-eDHFR(69K6)-HA-mNeonGreen<<kan | Fig. 4 | This study, L972 |
| AN033 | h- orc2-eDHFR(69K6)-HA-mNeonGreen<<kan | Fig. 4 | This study, L972 |
| AN034 | h- orc4-eDHFR(69K6)-HA-mNeonGreen<<kan | Fig. 4 | This study, L972 |
| AN035 | h- cdc45-eDHFR(69K6)-HA-mNeonGreen<<kan | Fig. 4 | This study, L972 |
| AN036 | h- plo1-eDHFR(69K6)-HA-mNeonGreen<<kan | Fig. 4 | This study, L972 |
| AN217 | h- pom1-eDHFR(69K6)-HA-mNeonGreen<<kan | Fig. 4 | This study, L972 |
| AN218 | h- spg1-eDHFR(69K6)-HA-mNeonGreen<<kan | Fig. 4 | This study, L972 |
| AN219 | h- ark1-eDHFR(69K6)-HA-mNeonGreen<<kan | Fig. 4 | This study, L972 |
| AN220 | h- mal3-eDHFR(69K6)-HA-mNeonGreen<<kan | Fig. 4 | This study, L972 |
| AN221 | h- sfi1-eDHFR(69K6)-HA-mNeonGreen<<kan | Fig. 4 | This study, L972 |
| AN222 | h- cut2-eDHFR(69K6)-HA-mNeonGreen<<kan | Fig. 4 | This study, L972 |
| AN223 | h- suc1-eDHFR(69K6)-HA-mNeonGreen<<kan | Fig. 4 | This study, L972 |
| AN231 | h- alp7-eDHFR(69K6)-HA-mNeonGreen<<kan | Fig. 4 | This study, L972 |
| AN232 | h- alp14-eDHFR(69K6)-HA-mNeonGreen<<kan | Fig. 4 | This study, L972 |

*nat: nourseothricin resistance gene (natMX6).

*kan: kanamycin resistance gene (kanMX6).

*mNeonGreen: the codon-optimized mNeonGreen for *Schizosaccharomyces pombe*.

*eDHFR(69K6): $^{iK6}$DHFR.

#### Supplementary Methods

**General materials and methods.**

All chemical reagents and solvents were purchased from commercial suppliers (Watanabe Chemical Industries, Tokyo Chemical Industry, FUJIFILM Wako Pure Chemical Corp., and Kanto Chemical) and used without further purification. Reversed-phase HPLC was performed on a Hitachi LaChrom Elite system with UV detection at 220 nm using a YMC-Pack C4 column (10 × 250 mm or 20 × 250 mm). $^{1}$H NMR spectra were recorded on a Bruker AVANCE III HD400SJ (400 MHz) spectrometer. $^{1}$H NMR chemical shifts were referenced to tetramethylsilane (0 ppm). High-resolution mass spectra were measured on a Thermo Scientific Exactive Plus Orbitrap mass spectrometer.

**Reagent abbreviations**

DBU: 1,8-diazabicyclo[5.4.0]-7-undecene
DIPEA: *N*,*N*-diisopropylethylamine
DMF: *N*,*N*-dimethylformamide
Fmoc-Adox-OH: Fmoc-8-amino-3,6-dioxaoctanoic acid
HATU: 1-((dimethylamino)(dimethyliminio)methyl)-1*H*-[1,2,3]triazolo[4,5-*b*]pyridine 3-oxide hexafluorophosphate
HBTU: 1-((dimethylamino)(dimethyliminio)methyl)-1*H*-benzo[*d*][1,2,3]triazole 3-oxide hexafluorophosphate
HOAt: 3*H*-[1,2,3]triazolo[4,5-*b*]pyridin-3-ol
HOBt: 1*H*-benzo[*d*][1,2,3]triazol-1-ol monohydrate
NMP: *N*-methyl-2-pyrrolidone
*o*-Ns-Cl: 2-nitrobenzenesulfonyl chloride
TFA: trifluoroacetic acid
TIPS: triisopropylsilane

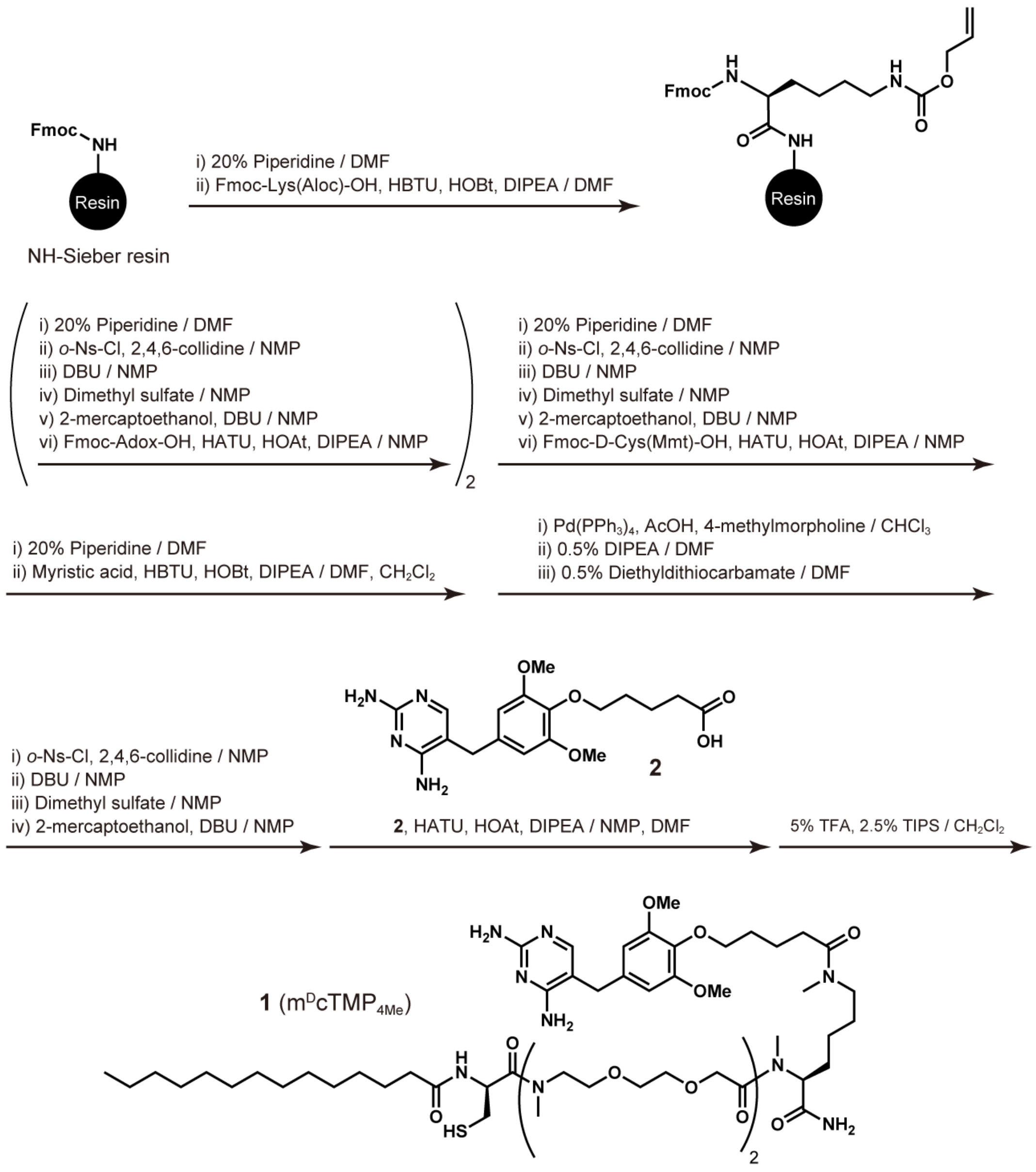


**Scheme S1. Synthesis of compound 1 (m$^{D}$cTMP$_{4Me}$)**

**Synthesis of compound 1 (m$^{D}$cTMP$_{4Me}$).**

Compound **1** was synthesized manually on Sieber amide resin (0.79 mmol/g) (38 mg, 30 μmol) using standard Fmoc-based solid-phase peptide synthesis protocols, as shown in **Scheme S1**. Fmoc deprotection was performed with 20% piperidine in DMF at room temperature for 15 min. After washing the resin with DMF, Fmoc-Lys(Aloc)-OH was coupled to the resin at room temperature with a mixture of Fmoc-Lys(Aloc)-OH (3.1 eq.), HBTU (3.0 eq.), HOBt (3.0 eq.), and DIPEA (6.0 eq.) in DMF. The coupling reaction was monitored by the Kaiser test [1]. After Fmoc deprotection and washing,

*N*-methylation and subsequent amino acid coupling reactions were performed using the procedure previously reported [2]. The Fmoc-deprotected resin was treated with a mixture of *o*-Ns-Cl (4.0 eq.) and 2,4,6-collidine (10.0 eq.) in NMP for 15 min. After washing the resin with NMP, the resin was treated twice with dimethyl sulfate (10.0 eq.) in NMP for 2 min each time. After washing the resin with NMP, the resin was treated twice with a mixture of 2-mercaptoethanol (10.0 eq.) and DBU (5.0 eq.) in NMP for 5 min each time, yielding the *N*-methylated *N*-terminus. Fmoc-Adox-OH was coupled to the resin using Fmoc-Adox-OH (3.1 eq.), HATU (3.0 eq.), HOAt (3.0 eq.), and DIPEA (6.0 eq.) in NMP. The coupling reaction was monitored by the chloranil test [3]. Subsequent Fmoc deprotection, *N*-methylation, and coupling reactions of Fmoc-Adox-OH and Fmoc-D-Cys(Mmt)-OH were performed as described above. After Fmoc deprotection, the *N*-terminus was myristoylated with a mixture of myristic acid (4.1 eq.), HBTU (4.0 eq.), HOBt (4.0 eq.), and DIPEA (8.0 eq.) in DMF/$CH_2Cl_2$ (1/1). After washing the resin with DMF, MeOH, and $CH_2Cl_2$, the resin was dried *in vacuo*. The Aloc group of the lysine residue was deprotected by treatment with $CHCl_3$ containing $Pd(PPh_3)_4$ (3.0 eq.), 5.0% acetic acid, and 2.5% 4-methylmorpholine under an Ar atmosphere. The resin was washed with DMF containing 0.5% DIPEA (×5), DMF containing 0.5% sodium diethyldithiocarbamate (×5), and DMF only (×5). The deprotected side chain of the lysine residue was then *N*-methylated as described above, and compound **2** [4] was coupled to the lysine side chain with a mixture of **2** (4.1 eq.), HATU (4.0 eq.), HOAt (4.0 eq.), and DIPEA (8.0 eq.) in NMP/DMF (1/1). After washing the resin with NMP, DMF, MeOH, and $CH_2Cl_2$, the resin was dried *in vacuo*. Deprotection and cleavage from the resin were performed with $CH_2Cl_2$ containing 5% TFA and 2.5% TIPS. The crude product was purified by reversed-phase HPLC using a semi-preparative C4 column (a linear gradient of MeCN containing 0.1% TFA and 0.1% aqueous TFA) to afford **1** as a white solid [8.4 mg, 22% (as a mono-TFA salt)].

$^1$H NMR (400 MHz, $CD_3OD$): mixture of rotamers δ 7.21 (1H, s), 6.56 (2H, s), 5.09–4.87 (2H, m), 4.36–4.18 (4H, m), 3.93 (2H, t, *J* = 6.0 Hz), 3.80 (6H, s), 3.76–3.49 (14H, m), 3.49–3.33 (4H, m), 3.21 (2H, m), 3.04 (3H, m), 3.00–2.75 (9H, m), 2.67 (2H, m), 2.47 (2H, m), 2.23 (2H, t, *J* = 4.0 Hz), 2.05–1.48 (10H, m), 1.38–1.15 (22H, m), 0.89 (3H, t, *J* = 6.0 Hz).
HRMS (ESI): calculated for $[M+H]^+$, 1163.7108; found, 1163.7061.

**Synthesis of m$^D$cTMP and hoeTMP**

m$^D$cTMP and hoeTMP were synthesized as described previously [5,6].
